# GDNF/RET signaling pathway activation eliminates Lewy Body pathology in midbrain dopamine neurons

**DOI:** 10.1101/752899

**Authors:** Piotr Chmielarz, Şafak Er, Julia Konovalova, Laura Bandrés, Irena Hlushchuk, Katrina Albert, Anne Panhelainen, Kelvin Luk, Mikko Airavaara, Andrii Domanskyi

## Abstract

Neurodegenerative diseases are associated with proteostasis disturbances and accumulation of fibrillar proteins into insoluble aggregates. Progressive age-related degeneration of dopamine neurons is a primary cause of motor dysfunctions in Parkinson’s disease (PD) and substantial evidence supports critical involvement of α-synuclein (α-syn) in the etiology of PD. α-syn is a cytosolic protein present in high concentrations in pre-synaptic neuronal terminals and a primary constituent of intracellular protein aggregates known as Lewy Neurites or Lewy Bodies. Progression of Lewy pathology is a characteristic feature in the PD brains caused by the prion-like self-templating properties of misfolded α-syn. Modelling Lewy pathology progression with application of exogenously prepared α-syn preformed fibrils, we discovered that glial cell line-derived neurotrophic factor (GDNF) prevented formation of α-syn aggregates in dopamine neurons in culture and *in vivo* after viral vector expression of GDNF. These effects were abolished by CRISPR/Cas9-mediated deletion of receptor tyrosine kinase *Ret*, the major GDNF signaling pathway. Similar to GDNF, expression of mutated constitutively active RET (RET_MEN2B) was able to protect dopamine neurons. GDNF protection against α-syn pathology progression was abolished by Src and attenuated by Akt pathway inhibitors. For the first time, we have shown the neurotrophic factor-mediated protection against the misfolded α-syn propagation in dopamine neurons, uncovered underlying receptor and intracellular signaling pathways. These results for the first time demonstrate that activation of GDNF/RET signaling can be an effective therapeutic approach to prevent Lewy pathology spread at early stages of PD.

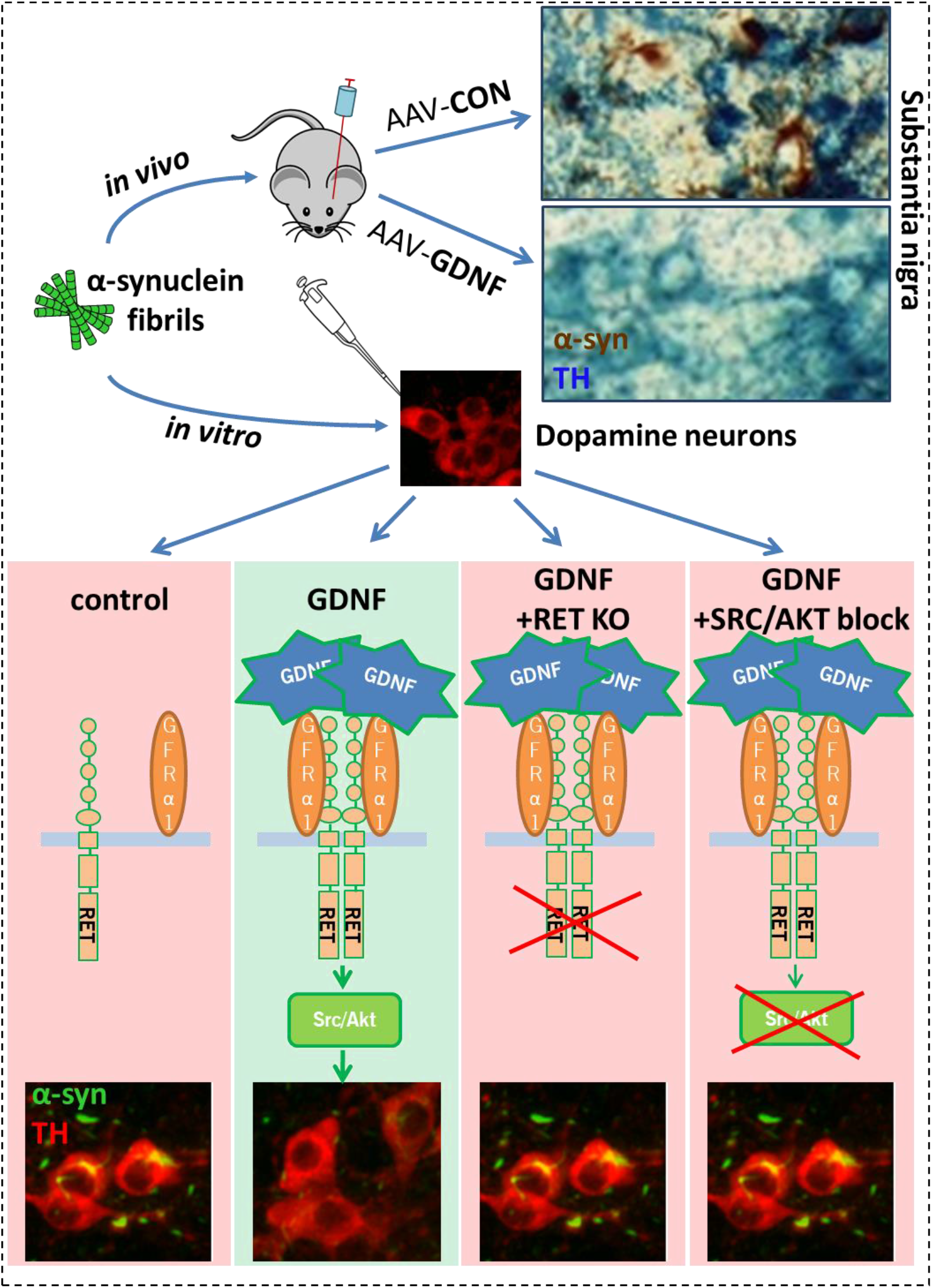

## Introduction

The presence of Lewy Neurites and Lewy Bodies – intraneuronal inclusions containing aggregated and phosphorylated (at serine 129; S129) α-syn as a main component – is a characteristic feature in the brains of both sporadic and familial Parkinson’s Disease (PD) patients (Anderson et al., 2006; Spillantini et al., 1997; Wong and Krainc, 2017).

Progression of Lewy pathology is evident already at the early stage of PD and coincides with compromised neuronal functions putatively by causing synaptic and mitochondrial dysfunction, cell stress, and deregulation of protein degradation pathways, which lead to neurodegeneration and disability in PD patients (Braak et al., 2003; Lashuel et al., 2013; Venda et al., 2010; Wong and Krainc, 2017). As α-syn aggregation is believed to be one of the earliest events in PD progression, treatments attenuating α-syn, aggregation and pathology spread should be applied to the patients at the earliest possible PD stages. Conversely, for late stage PD patients, transplantation of midbrain neural progenitors holds a great promise for cure (Barker et al., 2018). However, neurons transplanted into the brains of PD patients can accumulate Lewy Bodies (LBs), presumably, through transmission of misfolded prion-like proteins present in the PD patient’s brain (Kordower et al., 2008; Li et al., 2008). Therefore, treatments attenuating α-syn phosphorylation, aggregation and spread could be valuable also for late stage PD patients after transplantation therapy.

Glial cell line-derived neurotrophic factor (GDNF) promotes survival of mature dopamine (DA) neurons *in vitro* and *in vivo* (Kopra et al., 2017; Lin et al., 1993; Moore et al., 1996). GDNF binds and transduces signal through multiple receptors: RET tyrosine kinase (Durbec et al., 1996), Neuronal Cell Adhesion Molecule (NCAM) (Paratcha et al., 2003), syndecan-3 (Bespalov et al., 2011), N-cadherin and integrins (Kramer and Liss, 2015). The canonical receptor of GDNF is RET, a receptor tyrosine kinase family member, and its co-receptor, GDNF receptor alpha1 (GFRα1) that enhances GDNF affinity of RET (Durbec et al., 1996; Jing et al., 1996). GDNF had been extensively studied for its ability to rescue DA neurons from neurotoxic insults in multiple animal models (Paul and Sullivan, 2019). In comparison, there are relatively few studies addressing the effect of GDNF/RET signaling on the progression of Lewy pathology or α-syn aggregation (Decressac et al., 2011; Lo Bianco et al., 2004).

To model Lewy pathology progression, we utilized α-syn pre-formed fibrils (PFFs) that induce phosphorylation and aggregation of α-syn in cultured neurons and *in vivo* in the adult mice (Luk et al., 2012; Volpicelli-Daley et al., 2011). *In vivo* injections of PFFs recapitulate progression of α-syn aggregation and, in some mouse and rat strains, also neurodegeneration (Abdelmotilib et al., 2017; Paumier et al., 2015). This model is therefore well suited for the development of protective strategies to prevent α-syn aggregation and spread. Clarification of GDNF effectiveness against α-syn related pathology, in a model that is more representative of early stage PD, is of great clinical interest considering recent results of GDNF clinical trials, which clearly demonstrated feasibility and safety of long-term intracerebral GDNF administration, with good putamen coverage, minimal side effects and no anti-GDNF antibodies detected (Whone et al., 2019a; Whone et al., 2019b). Nonetheless, none of the double blind GDNF clinical trials have met clinical endpoints, as most GDNF treated patients had only modest improvements (Paul and Sullivan, 2019; Whone et al., 2019a; Whone et al., 2019b). The trials, however, included patients with moderate disease severity, at least 5 years (8 years on average) after initial motor symptoms, a stage at which DA cell loss is advanced and LB pathology is widespread (Kordower et al., 2013).

In the current study we demonstrate that GDNF can effectively protect DA neurons from progressing α-syn aggregation in models of early Lewy pathology and uncover underlying signaling pathways.

## Results

### PFFs induce robust α-syn aggregation in cultured dopamine neurons

GDNF is one of the most potent survival promoting factors for DA neurons; however, its effect on α-syn aggregation has been controversial. To address it, we plated and maintained embryonic midbrain cultures in a medium lacking GDNF. In these culture conditions, application of α-syn PFFs to DA neurons led to the progressive appearance of clearly detectable aggregates of phosphorylated (at Ser129) α-syn (pαSyn) both in neuronal soma and in neurites of TH-positive neurons. Aggregates could be observed starting from around 3 days post PFFs, as small puncta, putatively in neurites. The amount of these small pαSyn-positive puncta greatly increased at 5 days post PFFs. At 7 days post PFFs, large LB-like aggregates in cell soma were observed (Fig. 1A).

**Figure 1.**
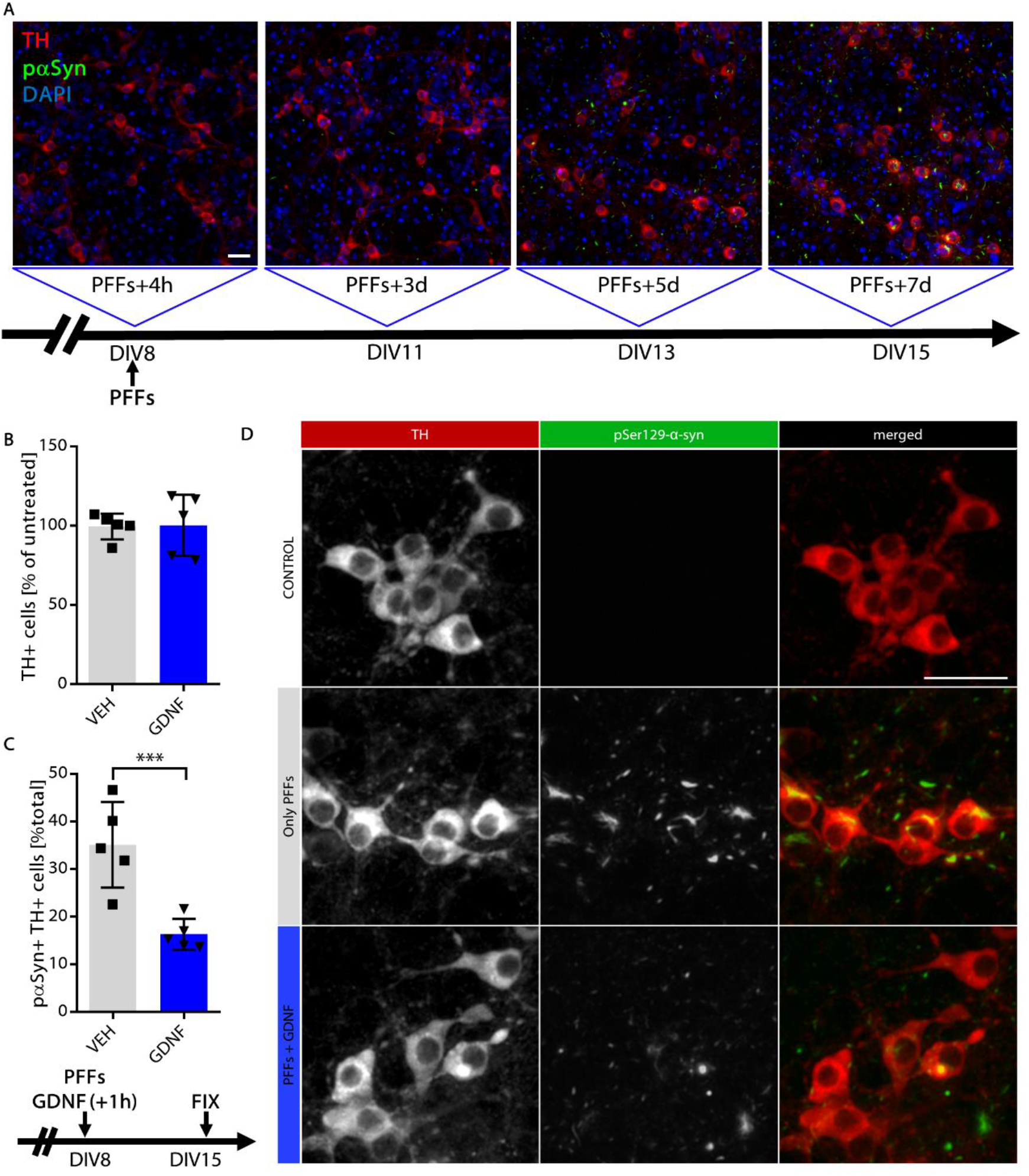
GDNF reduces accumulation of α-synuclein aggregates in cultured dopamine neurons. **(A)** Selected images depicting progressive accumulation of phosphorylated (at Ser129) α-synuclein (pαSyn) in midbrain neuronal cultures. α-synuclein aggregation was seeded with pre-formed fibrils (PFFs) at Day In Vitro (DIV) 8 and assessed by immunofluorescent staining on cultures fixed 4h, 3d, 5d and 7d after seeding. First pαSyn aggregates (green) can be seen at DIV11 outside of dopamine neurons’ soma (immunostained with anti-tyrosine hydroxylase (TH, red) antibody). Large intrasomal aggregates of pαSyn in fraction of dopamine neurons can be seen at DIV15 (7 days post PFF addition). **(B)** Neither PFFs no GDNF treatment (started at DIV8) affected dopamine neuron survival. **(C)** GDNF added 1h after PFFs significantly reduced number of dopamine neurons harboring Lewy Body-like intrasomal pαSyn aggregates (ratio paired t test). **(D)** Selected images of control (top) and PFF treated cells at DIV15 (7 days post PFF addition) from vehicle (middle) or GDNF treated (bottom) groups. ***p<0.001, n=5 independent experiments. Data are mean ±SD. Scale bars, 50 µm.

### GDNF reduces accumulation of α-syn aggregates in cultured dopamine neurons

Consistent with literature, omitting GDNF from the culture medium resulted in increased cell loss during first few days of culture (Planken et al., 2010), but most of the remaining cells survived for at least 15 days; therefore, number of DA cells was not further modulated by day *in vitro* (DIV) 8 addition of GDNF nor PFFs (Fig. 1B). Interestingly, addition of recombinant GDNF protein to neuronal cultures reduced the formation of pαSyn-positive LB-like aggregates in primary DA neurons (Fig. 1C, D **and Fig. S1**) but not in cultured hippocampal neurons (**Fig. S2**).

Similarly, lentiviral vector-mediated overexpression of GDNF under neuron specific human synapsin (hSyn) promoter (Domanskyi et al., 2015) was effective at reducing progression of α-syn aggregation assessed by immunostaining with antibodies detecting misfolded (Fig. 2A, B) or phosphorylated α-syn (Fig. 2D). Also, neither GFP, nor GDNF overexpression affected neuronal survival (Fig 2C, E). Importantly, addition of GDNF expressing lentiviral vectors after PFFs was also effective at attenuating α-syn aggregation (Fig 2D). These results suggest that GDNF activated intracellular signaling pathways preventing PFF-induced α-syn aggregation, rather than inhibiting phosphorylation of Ser129 or physically interacting with PFFs and/or competing for biding putative receptors for their cellular uptake.

**Figure 2.**
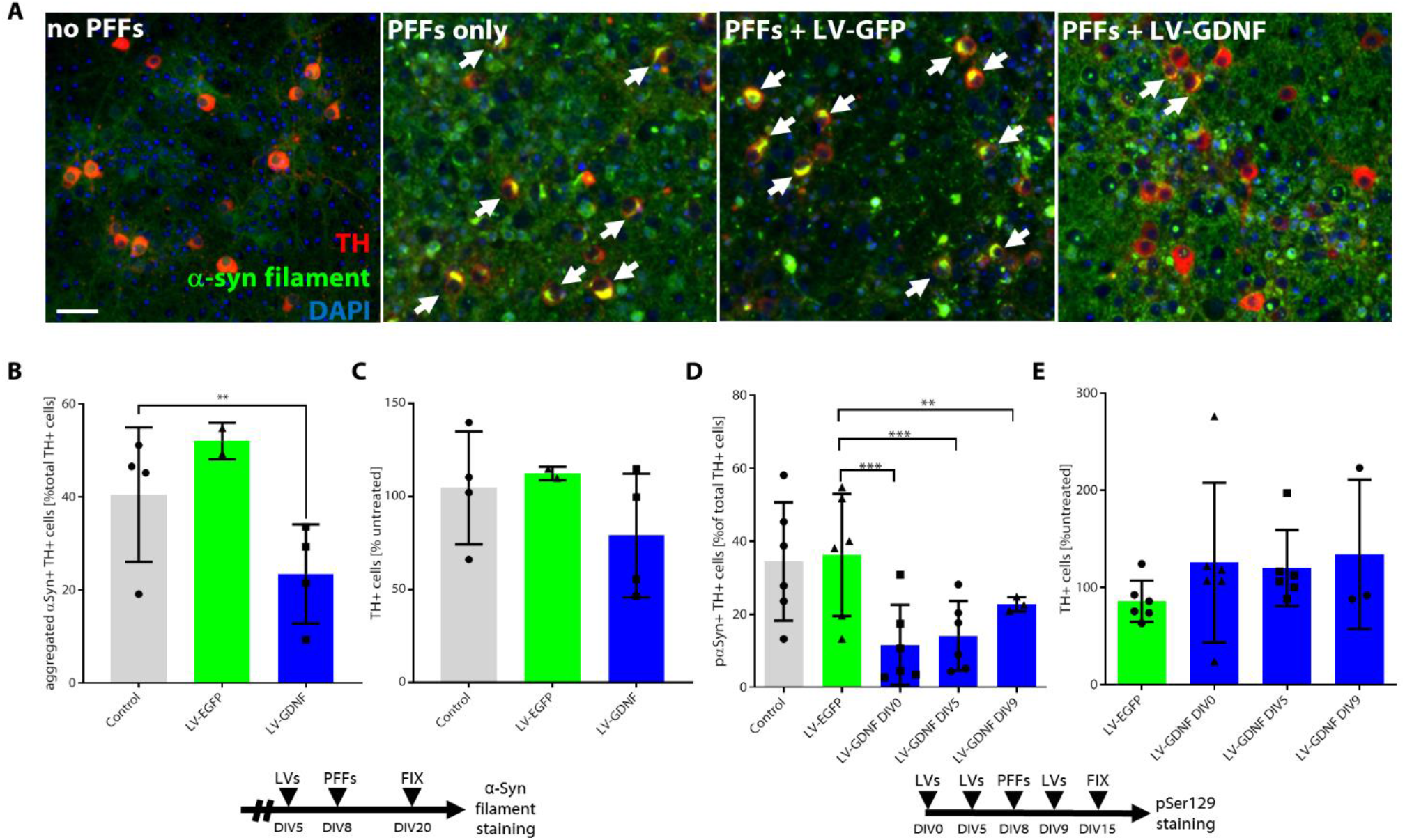
Lentivirus-mediated overexpression of GDNF reduces accumulation of pαSyn and misfolded α-syn in cultured dopamine neurons. **(A)** Compared to control (PFFs only) and LV-GFP (control vector), LV-GDNF reduces number of dopamine cells with misfolded α-syn, detected by α-syn filament antibody. **(B-E)** All the groups shown at graphs had PFF treatment on DIV8. **(B)** Quantification of TH-positive cells with aggregated α-syn shows that the number of cells with aggregates were significantly lower at the group transduced with LV-GDNF, 3 days before PFFs (ratio paired t test). **(C)** Dopamine cell survival assessed by number of TH-positive cells was not affected by either lentivirus vector. **(D)** LV-GDNF added 8 or 3 days before, or 1 day after PFFs significantly reduces number of dopamine cells with pαSyn in their soma (mixed effect ANOVA, Holm-Sidak’s multiple comparison test). **(E)** Dopamine cell survival assessed by number of TH-positive cells was not affected by either vector or different time points of transduction. **p<0.01, ***p<0.001, n=2-6 independent experiments. Data are mean ±SD. Scale bar, 50 µm.

### Virally mediated expression of GDNF in the substantia nigra prevents accumulation of Lewy Body-like aggregates of phosphorylated α-syn in DA neurons

To study the *in vivo* effect of GDNF, we used AAV-hGDNF to express it in the SN, and then utilized stereotaxic injections of PFFs to the mouse brain for the induction of *in vivo* α-syn aggregation (Fig. 3A) (Luk et al., 2012). Similar to the results obtained in primary midbrain cultures, GDNF strongly inhibited the accumulation of pαSyn positive aggregates in the SN (Fig. 3B, D, E), while the amount of LB and LN -like aggregates were similar in the cortex, striatum and amygdala – the regions not targeted by AAV-hGDNF vector (Fig. 3C). In some animals, GDNF overexpression also resulted in decreased intensity of TH staining, probably resulting from downregulation of TH expression as shown in literature (Chtarto et al., 2016; Georgievska et al., 2004) (**Fig. S3**).

**Figure 3.**
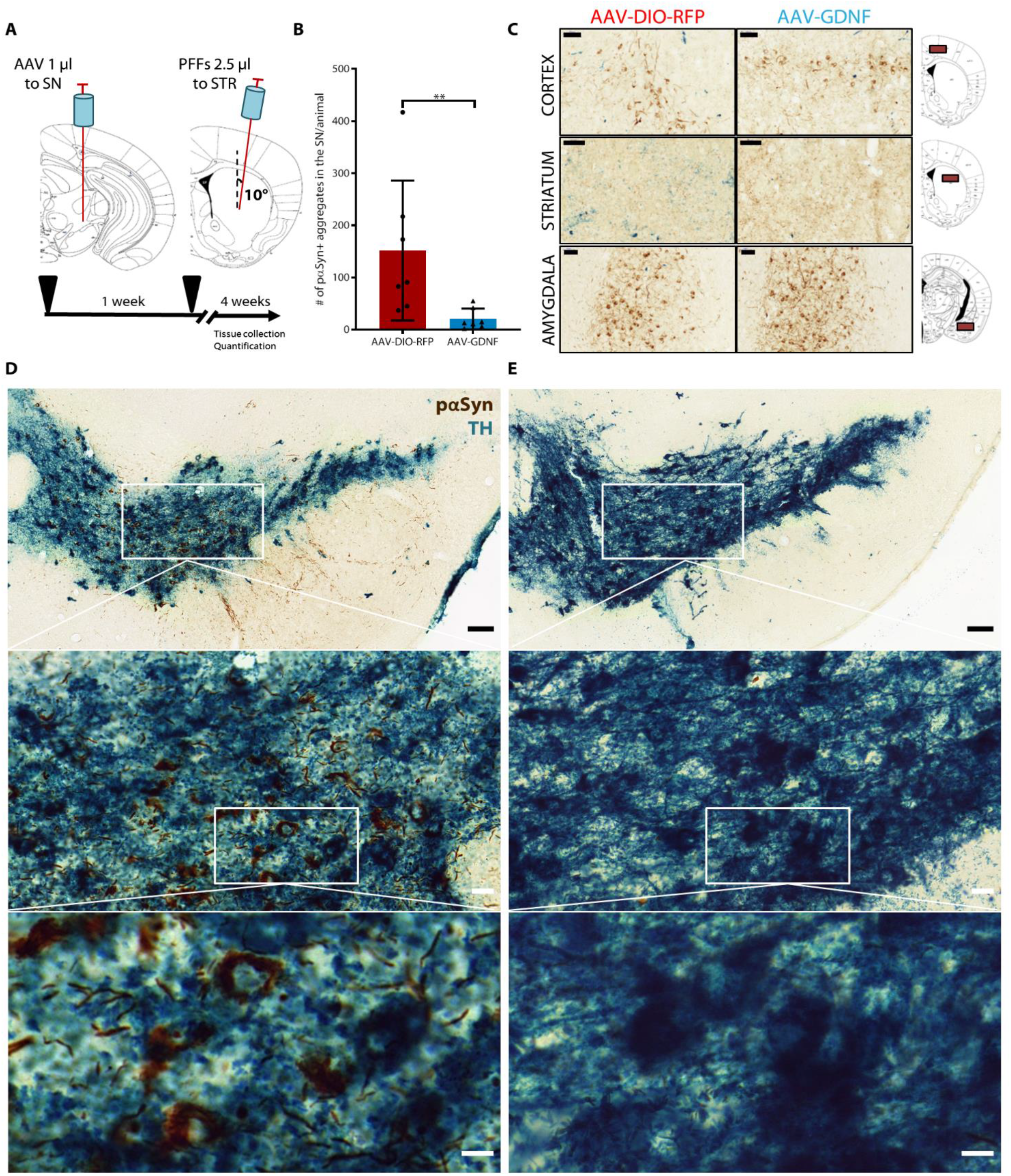
AAV-mediated overexpression of GDNF *in vivo* prevents accumulation of α-synuclein aggregates in the substantia nigra. **(A)** AAV-DIO-RFP (control) or AAV-GDNF was injected to the substantia nigra (SN), unilaterally. A week later, PFFs were injected to the striatum (STR), at the same side of the brain, and the brains were collected four week afterwards. **(B)** GDNF overexpression in the SN significantly reduces number of LB-like aggregates assessed by immunostaining for mouse pαSyn (Mann-Whitney test). **(C)** GDNF overexpression in the SN had only local effect and did not affect other brain regions (cortex, striatum, amygdala) in which pαSyn positive aggregates were detected. Scale bars, 50 µm. **(D)** Selected images depicting progressive accumulation of mouse pαSyn in TH-positive cells in the SN and **(E)** reduction in the number of pαSyn positive aggregates after GDNF overexpression. Scale bars, 100 µm, 20 µm and 10 µm, respectively. **p<0.01, n=7 animals (7-9 sections per animal). Data are mean ±SD.

### Effect of GDNF on accumulation of phosporylated α-syn is mediated by RET receptor

GDNF binds a co-receptor GFRα1 and activates receptor tyrosine kinase RET (Durbec et al., 1996; Jing et al., 1996; Kramer and Liss, 2015; Treanor et al., 1996; Trupp et al., 1996). It can also signal through other receptors, such as NCAM (Paratcha et al., 2003) and syndecan-3 (Kramer and Liss, 2015). To identify the cellular receptor mediating GDNF action on α-syn aggregation, we have utilized CRISPR/Cas9 genome editing to selectively target *Ret* in primary DA neurons (Fig 4A-D). CRISPR/Cas9-mediated deletion of *Ret* abolished GDNF effects on reducing accumulation of pαSyn aggregates (Fig. 4A, C).

**Figure 4.**
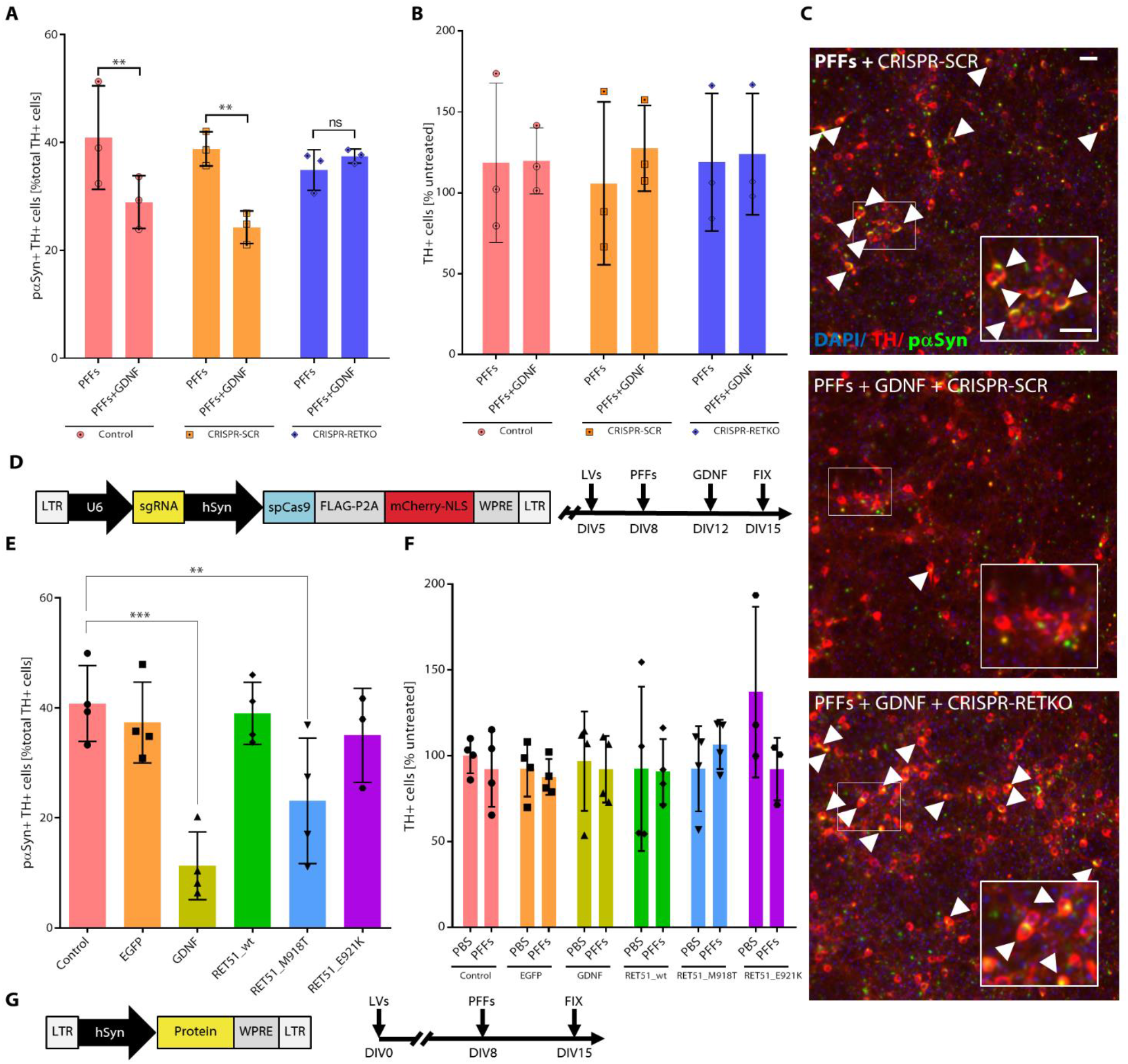
RET is essential for GDNF-dependent reduction of pαSyn accumulation in dopamine neurons. **(A)** Compared to control groups (no LVs and lentiCRISPR with scrambled (SCR) sgRNA), delivery of lentiCRISPR with sgRNA targeting *Ret* completely abolished the effect of GDNF protein on pαSyn aggregation (mixed model ANOVA and Holm-Sidak’s multiple comparison test). **(B, F)** Dopamine cell survival assessed by number of TH-positive cells was not affected by any of the treatments or lentiCRISPR vectors. **(C)** Selected images depicting progressive accumulation of pαSyn in TH-positive cells (control, top), reduction of aggregates after GDNF protein treatment (positive control, middle), and abolishment of GDNF effect in the presence of CRISPR-Cas9 targeting *Ret* (down). **(D)** For each individual lentiCRISPR vector, *spCas9* and mCherry-NLS reporter were expressed under human *Synapsin* (hSyn) promoter and sgRNA under U6 promoter. Lentivirus vectors were added 3 days before PFFs and 7 days before GDNF protein. **(E)** Lentivirus-mediated overexpression of constitutively active RET mutant (RET51_M918T) reduced pαSyn aggregation in TH-positive cells (mixed model ANOVA and Holm-Sidak’s multiple comparison test). **(G)** EGFP (control), GNDF, RET_wt RET51_M918T, RET_E921K (kinase dead mutant) proteins were delivered to dopamine neurons by lentivirus vector transduction and overexpressed under hSyn promoter, 8 days prior to addition of PFFs. *p<0.05, **p<0.01, ***p<0.001, n=3-5 independent experiments. Data are mean ±SD. Scale bars, 50 µm.

Conversely, GDNF effect on DA neurons could be mimicked by the lentivirus-mediated expression of constitutively active RET (RET_M918T, also known as MEN2B) (Mijatovic et al., 2007; Mijatovic et al., 2011), but not by overexpression of WT nor “kinase dead” RET_E921K mutant (Fig 4E-G). Neither *Ret* deletion nor overexpression affected neuronal survival (Fig 4B, F).

### Action of GDNF on α-syn accumulation involves activation of Src and Akt kinases

Using specific kinase inhibitors, we have found that Src kinase inhibitor SU6656 blocked the GDNF-mediated attenuation of α-syn aggregation, whereas MAPK pathway inhibitor U0126 had no effect (Fig. 5A). PI3K/Akt pathway inhibitor LY294002 increased the amount of DA neurons with LB-like aggregates independently from GDNF effect, which could be partially attenuated by the GDNF treatment (Fig. 5A). Neither SU6656, U0126 nor LY294002 caused significant effects on cell survival (Fig. 5B). Similarly, increase in basal number of pαSyn-positive cells was observed after treatment with highly specific Akt inhibitor MK2206, which also attenuated GDNF-induced reduction of pαSyn aggregation (Fig. 5C). Already at lowest tested concentration, MK2206 had detrimental effect on DA neuron survival, which was even more pronounced at higher concentrations and could be partially rescued by GDNF (Fig. 5D). All drug treatments followed the same schedule (Fig. 5E). A proposed simplified cascade of signaling events leading to reduced pathological α-syn accumulation in DA neurons upon RET activation is depicted in Fig. 5F.

**Figure 5.**
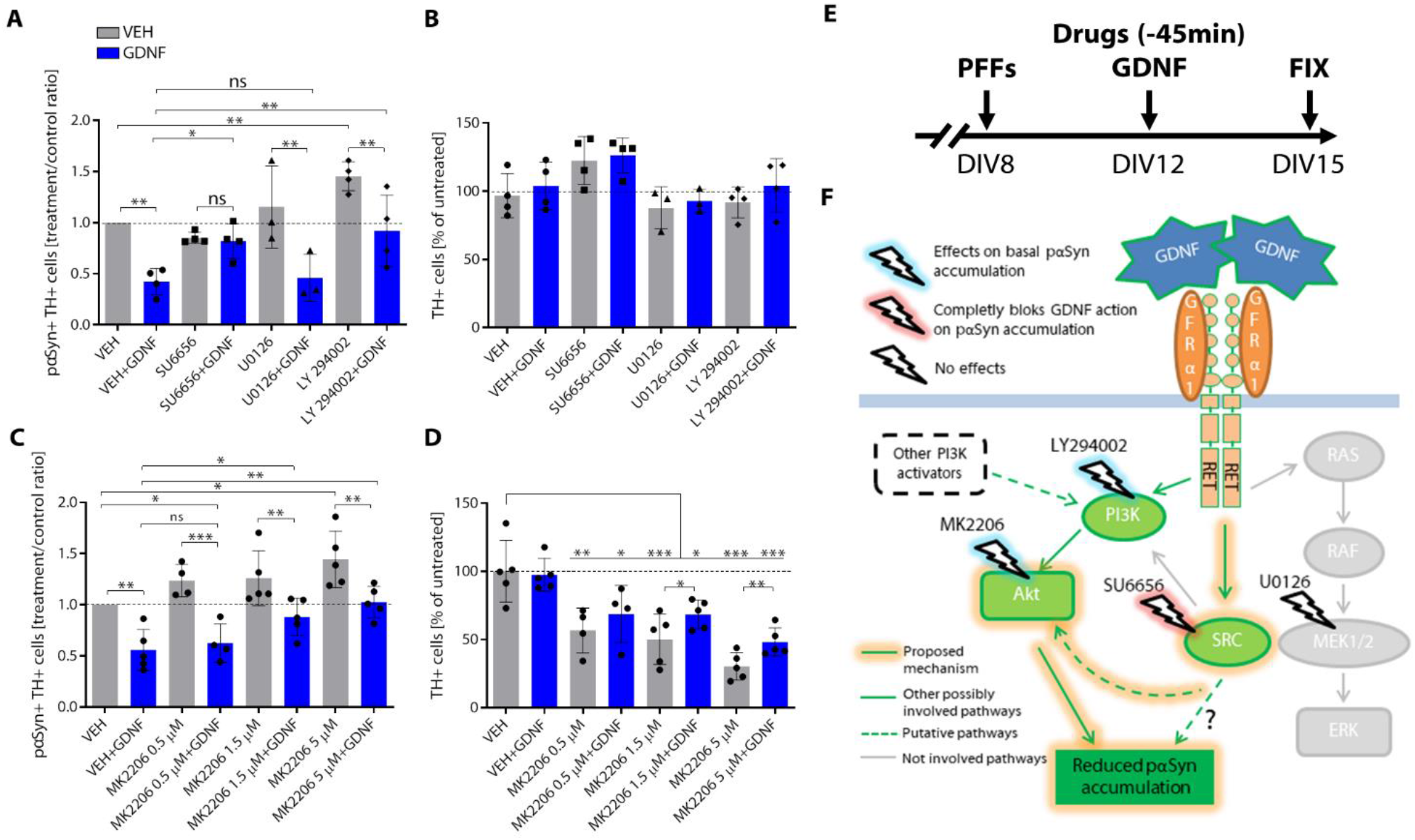
GDNF/RET effects on α-syn accumulation require Src and Akt activation. Dopamine neurons exposed to PFFs at Day In Vitro (DIV) 8 were treated at DIV12 with GDNF alone or after 45 minutes pretreatment with SRC, PI3K and MEK1/2, Akt inhibitors. **(A)** Quantification of large intrasomal aggregates of pαSyn in dopamine (TH-positive) cells. GDNF caused similar reduction of pαSyn aggregates in TH-positive cells when administered 4 days after PFFs as in previous experiments. Treatment with SRC inhibitor SU6656 abolished the effects of GDNF, while PI3K inhibition with LY294002 increased number of pαSyn positive dopamine cells; however, it did not abolish GDNF effect. Addition of U0126, a MEK1/2 inhibitor, did not affected GDNF effect on pαSyn accumulation in TH-positive cells (mixed effect ANOVA and Holm-Sidak’s multiple comparison test or one sample t test). **(B)** Dopamine cell survival assessed by number of TH-positive cells was not significantly affected by treatments. **(C)** MK2206, a direct allosteric Akt inhibitor attenuated GDNF effect on pαSyn accumulation in TH-positive cells (mixed effect ANOVA and Holm-Sidak’s multiple comparison test or one sample t test). **(D)** MK2206 also negatively affected dopamine cell survival which could be partially rescued by GDNF (mixed effect ANOVA and Holm-Sidak’s multiple comparison test or one sample t test). **(E)** Schematic timeline for all pathway inhibitors experiments. **(F)** Simplified scheme of intracellular GDNF/RET signaling with highlighted elements mediating effects on pαSyn accumulation. Increase of pαSyn after blockade of PI3K demonstrates its role in early steps of pαSyn accumulation after PFF treatment. However, the effect of GDNF could only be partially blocked by inhibiting PI3K. In contrast, complete abolishment of GDNF effect after Src inhibition suggest a critical role of non-canonical direct Src-mediated Akt activation or Akt-independent pathways. *p<0.05, **p<0.01, ***p<0.001, n=3-5 independent experiments. Data are mean ±SD.

## Discussion

Our data for the first time demonstrate that GDNF can slow or even stop accumulation of α-syn in models recapitulating spreading of misfolded α-syn – a neuropathological hallmark of PD. Importance of our finding is that for the first time we demonstrate the connection between two major therapeutic approaches for PD – neuroprotection through stimulation of trophic pathways and slowing down the spread of Lewy pathology. Therapies based on either of aforementioned approaches had been and/or are planned to be tested in clinic (clinical trials ##NCT03100149, NCT03318523, NCT03716570, NCT02618941, NCT02267434, NCT03611569, NCT03272165, NCT01621581, NCT03295786, NCT01621581) (Huttunen and Saarma, 2019; Jankovic et al., 2018; Paul and Sullivan, 2019; Weihofen et al., 2019). Our data for the first time indicate a possibility to design neurotrophic signaling-based therapies with two-fold mode of action addressing both survival and Lewy pathology spread. However, such therapies might require modified clinical approaches e.g. treatment of early stage patients, which was already suggested as a way to increase efficiency of neurotrophic therapies for PD (Paul and Sullivan, 2019). Identification of such patients will be increasingly feasible with the advancement of diagnostic methods for the early detection of α-syn oligomers (e.g. in skin biopsy samples (Doppler et al., 2017) or from cerebrospinal fluid (Kang et al., 2019)).

Importantly, there is ongoing controversy if GDNF therapies can at all be effective in PD with Lewy pathology. Decressac et al. have claimed that in rat α-syn overexpression models and PD patients with Lewy pathology α-syn accumulation can lead to downregulation of *Ret*, which impairs GDNF receptor signaling thus rendering GDNF treatments ineffective (Decressac et al., 2012; Decressac et al., 2013; Decressac et al., 2011). However, later studies have not observed RET downregulation neither in PD animal models nor in post mortem PD patient brains (Su et al., 2017). In addition, PET imaging in recent GDNF clinical trials showed clear effects of GDNF in PD patients – i.e. increase in 18F-DOPA uptake (Whone et al., 2019a). While interpreting results of Decressac et al., it is important to note that they were done in models utilizing AAV-mediated overexpression of α-syn, which might have resulted in very high α-syn expression levels (Hoffer and Harvey, 2011). This model does not recapitulate spreading nature of Lewy pathology; however, might represent a late stage PD or some subpopulation of patients. In contrast, our studies utilized model of early, spreading Lewy pathology. Considering our finding that GDNF/RET signaling limits accumulation of misfolded α-syn, it is intriguing to speculate that after reaching certain threshold level, misfolded α-syn accumulation and impairment of GDNF/RET signaling might form a vicious cycle with misfolded α-syn impairing GDNF/RET action thus leading to even faster progression of Lewy pathology.

The progression of LB-like pathology, i.e. accumulation of aggregated, phosphorylated at Ser129 α-syn in our midbrain cultures, was similar to what was observed in other publications (Volpicelli-Daley et al., 2014; Volpicelli-Daley et al., 2011). In the time frame of our experiments, we did not observe any loss of DA neurons. Other studies also did not report DA cell loss at 7 days post-PFFs, and modest cell loss was observed after longer periods (i.e. 14 days post PFFs) and/or additional neurotoxic insult to the cells (Tapias et al., 2017; Volpicelli-Daley et al., 2014). Moreover, the lack of cell death at the early time points after induction of the pathological α-syn aggregation is consistent with reports showing that while the rodents injected with PFFs develop robust α-syn aggregation, such loss of DA neurons was observed only several months after PFF treatment. Similarly, the transgenic mice overexpressing α-syn exhibit DA neurodegeneration only at old age (Lee et al., 2002; Luk et al., 2012; Paumier et al., 2015; Richfield et al., 2002; Thiruchelvam et al., 2004). Accordingly, motor abnormalities in PFF-treated or α-syn overexpressing animals were observed after several months post-injection of PFFs or at advanced age. Despite years of research, clear cause of DA neurons’ loss in PD remains undefined, with support gained by the “multiple hit” hypothesis suggesting that neurodegeneration in PD is caused by a combination of several factors, one of which is α-syn misfolding (Surmeier et al., 2017). Moreover, recent research on human patient iPSC-derived DA neurons shows that they are more vulnerable than mouse DA neurons due to higher DA content and even then, robust pathological phenotypes are only observed after very long (several months) culturing periods and/or introduction of multiple detrimental factors, with α-syn-related pathology among them (Burbulla et al., 2017). Our data is in line with this research supporting evidence that aggregation of α-syn might be just one of multiple hits necessary for the demise of DA neurons in PD.

In our experiments, we demonstrated that GDNF is effective at reducing Lewy pathology progression at least at early time points after PFFs application. Efficacy of the GDNF treatment *in vivo* was much greater than that observed *in vitro*, leading to almost complete abolishment of LB-like aggregates in the SN of mice overexpressing GDNF at the region. This might be due to high levels of GDNF achieved *in vivo* by AAV-hGDNF expression, as we and others have previously observed (Kirik et al., 2000; Penttinen et al., 2018; Tenenbaum and Humbert-Claude, 2017). Another explanation is that the cultured cells were exposed to constant, high PFF concentration, which resulted in much faster acquisition of pathology, while *in vivo* experiments rely on uptake of PFFs by DA axons in the striatum and transport of PFFs into the cell body, resulting in much slower development of α-syn aggregates in DA neuron soma, which was more efficiently prevented by GDNF. Altogether, these data suggest that while in our *in vitro* setup GDNF is moderately effective, it reliably protected DA neurons *in vivo*, at least when administered before the development of LB pathology.

GDNF exerts its function through several receptors with partially overlapping effector pathways. Our data demonstrate that RET is indispensable for effects of GDNF in reducing pαSyn accumulation. Moreover, overexpression of different isoforms of RET shows that constitutively active RET mutant is sufficient to reduce pαSyn accumulation. Constitutively active RET receptor was previously shown to exert protective effects on DA neurons (Mijatovic et al., 2011); however, its ability to reduce progressive accumulation of pαSyn has never been studied before. Our findings strongly support the validity of RET as a target for disease-modifying therapy for PD. Importantly, the presented data supports introduction of RET activating therapy at early stages of the disease, both to exert pro-survival effects on DA neurons and to protect them from the development of Lewy pathology.

Effect of GDNF on pαSyn accumulation was completely abolished by inhibition of Src kinase but not by inhibition of MAP signaling pathway. Blockade of PI3K or direct blockade of Akt with a specific inhibitor resulted in increase of pαSyn accumulation, which could be partially rescued by the GDNF treatment. In our studies we used MK2206, a highly specific allosteric inhibitor of Akt, not competitive to ATP binding, which was shown to effectively block canonical PI3K/Akt pathway, by preventing T308 and S473 Akt phosphorylation (Manning and Toker, 2017; Meuillet, 2011; Xing et al., 2019). However, MK2206 does not inhibit kinase domain of Akt directly and, therefore, does not preclude non-canonical activation.

Classically, Akt activation by GDNF/RET is thought to be mediated by PI3K signaling in either Src dependent or Src independent manner (Fayard et al., 2011; Kramer and Liss, 2015). Interestingly, Src is also able to activate Akt through non-canonical PI3K independent pathway by phosphorylating Akt on Y315 and Y326 residues (Mahajan and Mahajan, 2012), although the role and prevalence of this mechanism is poorly described in literature. In fact, mutated form of RET – RET/PTC (rearranged in transformation/papillary thyroid carcinomas) was shown to promote Y315 and Y326 Akt phosphorylation, presumably through Src activation (Jung et al., 2005). Moreover, Src can be involved in Akt activation by both PI3K-dependent and -independent pathways even when transducing signal from the same receptor (Lodeiro et al., 2011). While this pathway has been demonstrated in cancer cells, our data suggest such mechanism may also be involved in GDNF/RET dependent Akt activation in DA neurons.

Our results could be explained by PI3K-independent, Src-dependent activation of Akt following GDNF/RET activation. However, we cannot fully rule out that Akt and Src are independently affecting pαSyn accumulation in DA neurons. Therefore, our data warrant further studies of possible PI3K-independent, Src-dependent activation of Akt, or other Src-dependent Akt-independent pathways by GDNF and their role in protection against Lewy pathology. Due to limited availability of ligands for dissection of different Akt activation mechanisms, such studies should employ genetically modified Ret, Akt and/or Src. Blocking Akt with MK2206 also resulted in the loss of DA neurons; however, this neuronal loss was similar in control and PFF treated groups. Therefore, the effect was not dependent on pαSyn accumulation and did not impact the interpretation of obtained results.

Precise understanding of protective intracellular pathways activated by GDNF in DA neurons is of great importance for engaging RET as a target for PD therapy. For example, putative biased GDNF mimetics, activating Akt through PI3K-independent, Src-dependent pathway may offer an attractive therapy for early PD, protecting against Lewy-pathology, with decreased risks associated with strong Akt activation. Interest in developing GDNF or GDNF-mimetic therapy for PD had been declining in the past several years due to inconclusive results of clinical trials with GDNF (Paul and Sullivan, 2019) and preclinical and patient data suggesting the loss of RET receptor in later stages of PD (Decressac et al., 2012; Decressac et al., 2013; Decressac et al., 2011). However, as discussed above, this might not be true for all population of patients, especially those at earlier stages of the disease, and recent clinical trials for GDNF, while not meeting clinical endpoints, demonstrated feasibility of safe, long term GDNF administration (Whone et al., 2019a; Whone et al., 2019b). Importantly, discussing the modest efficacy of GDNF in the latest clinical studies, the authors suggested future modifications of GDNF trials to increase GDNF dosing or include patients at early stages of PD, what is strongly supported by our data. In addition, less invasive methods of activation of RET receptor together with emerging biomarkers for early PD could provide strategy to prevent disease progression. Actually, GDNF-mimetics are being actively developed (Ivanova et al., 2018; Sidorova et al., 2017) and our data strongly support their potential therapeutic application for the treatment of PD at early stages.

Precise mechanism by which activation of RET and downstream pathways reduces accumulation of aggregated α-syn remains unclear. It has been recently proposed that upon their endocytosis, PFFs remain for several days in the endosomal pathway, where they seem to undergo partial processing (Apetri et al., 2016; Flavin et al., 2017) and are subsequently released to the cytoplasm, then promoting fibrillization of endogenous α-syn protein (Bieri et al., 2018). Therefore, one can speculate that mechanism of RET/Akt action on α-syn could be either interfering with uptake of PFFs, modifying their early processing in endosomal pathway and/or release to cytoplasm, or enhancing clearance of misfolded cytoplasmic α-syn aggregates. Interfering with PFF uptake seems unlikely, as PFFs were shown to be uptaken very rapidly, within hours of application (Karpowicz et al., 2017), while we show that GDNF was equally effective when applied before and up to 4 days after PFF seeding. Interestingly approximately 5 days after their addition, PFFs were reported to appear in the cytoplasm in neuronal soma, recruiting endogenous α-syn into pSer129-αsyn positive aggregates (Apetri et al., 2016; Karpowicz et al., 2017). As inhibition of lysosomal processing was shown to accelerate release of PFFs to cytoplasm and recruitment of endogenous α-syn (Karpowicz et al., 2017), it is possible that RET/Akt activation is altering the processing of the endocytosed fibrils and decreasing their chance of initiating α-syn aggregation in cell soma. Alternatively, activation of GDNF/RET signaling could lead to enhanced clearance of α-syn aggregates, which, if started early when aggregates are few and small, could prevent snowballing increase of α-syn aggregation.

Altogether, our results, for the first time, demonstrate that GDNF/RET signaling can protect DA neurons from pathological pαSyn accumulation. Moreover, we show that treatment with neurotrophic factors, or neurotrophic factor mimetics activating the correct intracellular pathways might be effective specifically at the early stage of pathology. These results support early initiation of treatment with neurotrophic factors or neurotrophic factor mimetic in PD patients.

## Materials and Methods

### Animal experiments

63-week-old C57Bl/6NCrI mice (8 males, 6 females) were housed at a 12-hour light–dark cycle, with free access to water and food. All stereotaxic injections were performed under 4% isoflurane. AAVs (AAV1-DIO-iRFP or AAV-GDNF (Penttinen et al., 2018; Richie et al., 2017)) were injected to the SN (A/P −3.3 mm, M/L −1.2 mm, D/V −4.9 mm from bregma; at 10° angle). One week later, 5 µg of α-syn PFFs (2 µg/µL in PBS) were injected to the striatum (A/P 0.7 mm, M/L −2.2 mm, D/V −3.0 mm from bregma). After 4 weeks, the brains were collected after anesthetization of the animals with sodium pentobarbital (Orion Pharma, i.p. 90 mg/kg), followed by intracardial perfusion with PBS and 4% PFA. All animal experiments were approved by the Finnish National Board of Animal Experiments (license number: ESAVI/7812/04.10.07/2015) and were carried out according to the European legislation on the protection of animals used for scientific purposes.

### Immunohistochemistry of free-floating sections

The brains were sectioned as described in (Albert et al., 2019). Briefly, the brains were fixed with 4% PFA, put into a 20% sucrose solution, frozen in a cryostat (Leica CM3050), cut into 40-μm coronal sections covering the striatum (approximately 1.5 to −0.8 mm relative to bregma) and the midbrain (approximately −2.7 to −4.0 mm relative to bregma), and stored in 20% glycerol with 2% DMSO in PBS at −20°C until immunostaining. Every sixth section was used for immunostaining. The sections were thawed at room temperature (RT), rinsed with PBS and after 15 minutes blocking of endogenous peroxidase (0.3% H_2_O_2_ in 1:1 PBS:methanol), rinsed twice with 0.3% Triton X-100 in PBS (PBST) and blocked for 1 hour with 5% normal horse serum (NHS; Vector, S-2000) in 0.3% PBST. Sections were incubated with the first primary antibody (monoclonal rabbit anti-phosphoSer129-α-synuclein, Abcam, ab51253; 1:10000) in NHS blocking buffer overnight, at +4°C. Next, sections were rinsed and incubated with the secondary antibody (Vector Laboratories anti-rabbit or anti-mouse biotinylated secondary antibody, PK-4001 and PK-4002; 1:500) in NHS blocking buffer for 1 hour, at RT. After rinsing, sections were incubated in avidin-biotinylated horseradish peroxidase (ABC Kit, Vector Laboratories) in PBST for 20 minutes, rinsed, and developed with 3,3-diaminobenzidine-4 HCL (DAB) (DAB peroxidase substrate kit, SK-4100, Vector Laboratories) in water for 2-5 minutes, rinsed with PBS. Following 15 minutes 0.3% H_2_O_2_ blocking, sections were rinsed with PBST thrice. Then 15 minutes of Avidin and Biotin incubations (Vector Laboratories, SP-2001) were done. Sections were rinsed thrice and incubated in the second primary antibody (monoclonal mouse anti-TH, Millipore, MAB318; 1:3000) in NHS blocking buffer overnight at +4°C. Same secondary staining steps were repeated except developing, which was done with HistoGreen (HRP substrate kit, E109, Linaris) in water for 1-2 minutes. Sections were rinsed with water and placed on coated glass slides. Slides were allowed to dehydrate overnight at RT and mounted with Coverquick 2000 mounting medium. The slides were digitalized on Pannoramic P250 Flash II Whole Slide Scanner (3DHistech), with 20X magnification (extended focus, 0.22 μm/pixel resolution). The number of LB-like aggregates was counted manually, from 8-9 sections covering the SN. Immunohistochemistry of the sections and quantification of LB-like aggregates were performed by blinded experimenters.

### Primary embryonic midbrain cultures

Ventral midbrain floors were isolated from wild-type NMRI E13.5 mouse embryos, as described in (Planken et al., 2010). Cells were plated in microislands on 96-well plates and incubated in 5% CO2, at 37°C.

Recombinant mouse α-syn PFFs (5 mg/ml) were diluted 1:50 in PBS and sonicated at high power (10 cycles, 30 seconds on/ 30 seconds off) with Bioruptor sonicator (Diagenode, Liege, Belgium). On DIV8, 2.5 µg/mL of PFFs were added to cultures; 50 ng/ml recombinant GDNF protein (PeproTech, Rocky Hill, NJ, USA or Prospec, CYT-305) was added on DIV8 or DIV12. PBS was used as control.

For pathway analyses, 2 µM SU6656 (Src kinase inhibitor) (Tocris, 6475), 10 µM LY294002 (PI-3 kinase inhibitor) (Tocris, 1130), 10 µM U0126 (MEK1/2 inhibitor) (Tocris, 1144), 5 µM MK2206 (Akt kinase inhibitor) (Selleckchem, S1078) in DMSO were added on DIV12, ∼45 minutes before GDNF treatment. DMSO was used as vehicle control.

On DIV15 the cells were fixed 20 minutes at RT with 4% PFA and stored in PBS at +4°C until immunostaining. All experimental timelines are stated in corresponding figures.

### Immunohistochemistry of primary embryonic midbrain cultures

Fixed plates were rinsed with PBS twice, followed by 15 minutes of permeabilization with 0.2% Triton X-100 in PBS (PBST) and 1 hour of blocking with 5% NHS in PBST. The incubation with primary antibodies (monoclonal mouse anti-TH, Millipore, MAB318 or polyclonal sheep anti-TH, Millipore, ab1542; monoclonal rabbit anti-phosphoSer129-α-synuclein, Abcam, ab51253; rabbit monoclonal anti-Alpha-synuclein filament, Abcam, ab209538; all at 1:2000 dilution) in blocking solution was done at +4°C overnight. Thrice rinse with PBS was done before 1-hour incubation with secondary antibodies (donkey anti-mouse AlexaFluor 488, Thermo Fisher Scientific, A21202 or donkey anti-mouse AlexaFluor 568, Thermo Fisher Scientific, A10037; donkey anti-rabbit AlexaFluor 647, Thermo Fisher Scientific, A31573; 1:400), at RT. After PBS rinse, the cells were stained for 10 minutes with 200 ng/ml 4’,6-diamidino-2-phenylindole (DAPI). Plates were stored at +4°C until imaging.

Imaging of primary cells in 96-well black view plates (PerkinElmer, 6005182) was performed with CellInsight CX5 High Content Scanner (Thermo Fisher Scientific) or ImageXpress Nano Automated Imaging System (Molecular Devices), fitted with 10X objective. From each well 36 (CellInsight CX5) or 9 (ImageXpress Nano) view fields were acquired, encompassing entire microisland culture.

Quantification of TH-positive cells with or without LB-like pαsyn-positive aggregates was performed with CellProfiler and CellAnalyst software packages (Carpenter et al., 2006; Jones et al., 2009; Jones et al., 2008). Detailed CellProfiler image analysis pipelines can be provided upon request. Briefly, the first pipeline and CellProfilerAnalyst were used to quantify the numbers of TH-positive cells per well and subsequently classify identified TH-positive cells into LB-positive and LB-negative, utilizing supervised machine learning implemented in CellProfilerAnalyst.

### Statistical analysis

*In vitro* data which was performed in multiple independent experiments (plates) was analyzed utilizing randomized block ANOVA design, matching groups from different plates (Lew, 2007). Statistical significance was calculated by mixed model one-way or two-way analysis of variance (ANOVA) followed by Holm-Sidak’s multiple comparison test or one sample t test (vs ratio of control group. i.e. 1.0). Data from *in vivo* experiment was tested with Mann-Whitney nonparametric test. All statistical analyses were performed in GraphPad Prism 8.20 software (GraphPad Software, Inc). Data in text and figures are represented as means ± SD with individual data points plotted as overlay.

## List of Supplementary Materials

### Supplementary Materials and Methods

Primary embryonic hippocampal cultures

Cloning of lentiviral transfer vectors and CRISPR/Cas9 constructs

Production of lentivirus vectors

### Supplementary Figures

Fig S1. GDNF treatment up to 4 days after PFF addition reduces the accumulation of α-synuclein aggregates in cultured dopamine neurons.

Fig S2. GDNF does not prevent pαSyn accumulation in hippocampal neurons.

Fig S3. Accumulation of LB-like pSer129 α-synuclein aggregates in the SN of PFF-injected mice with or without GDNF overexpression.

## Acknowledgements

We thank Prof. Mart Saarma for critical comments and suggestions during implementation of this project, Congjun Zheng for establishing and culturing neuronal cells, Aastha Singh for experiments with hippocampal neurons, and the Light Microscopy Unit and HistoScanner service at the Institute of Biotechnology, University of Helsinki, for imaging immunostained cells and tissue sections. This work was supported by grants from 3i-Regeneration by Tekes (Finnish Funding Agency for Innovation), Academy of Finland #309489, #293392, #319195; and Sigrid Juselius Foundation.

## Author contributions

P.C. and Ş.E. designed and performed most of the experiments, analyzed the data and wrote the manuscript together with A.D.; J.K. and L.B helped with the experiments and data analysis; I.H. assisted with the *in vitro* experiments on dopamine cultures; K.A. designed and performed the animal experiments and proof-read the manuscript; A.P. helped with the design and execution of *in vitro* experiments with PFFs and hippocampal cultures; K.L. provided the PFFs and proof-read the manuscript; M.A. and A.D. supervised the research and provided critical feedback for the article.

## Data and materials availability

All data associated with this study are available in the main text or the supplementary materials.

