## Supplemental Materials and Figures for "GDNF/RET signaling pathway activation eliminates Lewy Body pathology in midbrain dopamine neurons"

**Supplementary Materials**

**Primary embryonic hippocampal cultures**

Hippocampi were isolated from wild-type NMRI E15-17 mouse embryos, as described in (Seibenhener and Wooten, 2012), at day *in vitro* (DIV) 0. Cells were plated on 96-well plates and incubated in 5% CO2, at 37°C. 2.5 µg/mL of recombinant mouse α-syn PFFs (prepared as described in the Materials and Methods) were added to cultures on DIV7. 50 ng/ml recombinant GDNF protein (PeproTech, Rocky Hill, NJ, USA or Prospec, CYT-305) or PBS (control) were added on DIV7 30 min before PFFs. On DIV14 the cells were fixed 20 min at room temperature with 4% paraformaldehyde (PFA) and stored in PBS at +4°C until immunostaining. For immunohistochemistry, mouse anti-NeuN, clone A60 (Millipore, MAB377, 1:400 dilution) and monoclonal rabbit anti-phosphoSer129-α-synuclein (Abcam, ab51253, 1:2000 dilution) were used as primary antibodies. Fixed plates were stained, imaged and analyzed similarly to the embryonic midbrain cultures, as described in the Materials and Methods.

**Cloning of lentiviral transfer vectors and CRISPR/Cas9 constructs**

A 452 nt fragment of human *Synapsin 1* (hSYN) gene promoter was PCR amplified using Phusion high fidelity DNA polymerase (Thermo Scientific) and the following primers, BcuI_hSYN_for 5'ATACTAGTAGTGCAAGTGGGTTTTAGGACC and EcoRI_hSYN_rev 5'TGGAATTCGACTGCGCTCTCAGG, digested with FastDigest-*Bcu*I/-*Eco*RI (Thermo Scientific) and cloned into the pCDH-CMV-T2A-EGFP vector (System Biosciences) digested with the same restriction enzymes, resulting in pCDH-hSYN-T2A-EGFP construct. To obtain the pCDH-hSYN-hGDNF lentivirus transfer plasmid, the coding sequence of pre-α-pro-GDNF was PCR amplified from human tissue cDNA with the following primers, GDNF_For 5'TTGGATCCATGAAGTTATGGGATGTCGT, GDNF_Rev GAGTCGACTCAGATACATCCACACCTTT, digested with FastDigest-BamHI/-SalI (Thermo Scientific) and cloned into a pCDH-hSYN-T2A-EGFP vector digested with the same enzymes. To obtain the pCDH-hSYN-hRET51wt, pCDH-hSYN-hRET51-M918T, and pCDH-hSYN-hRET51-E921K lentivirus transfer plasmids, the corresponding RET51 sequences were PCR amplified from mammalian expression vectors (gift from Prof. Mart Saarma) with the following primers, hRET_for 5'AATTTAAATCGGATCCGCCACCATGGCGAAGGCGACGTC, hRET_rev 5'GAGGTTGATTGTCGACTTAACTATCAAACGTGTCCATTAATTTTGC and cloned into a pCDH-hSYN-T2A-EGFP vector digested with FastDigest-*Bam*HI/-*Sal*I (Thermo Scientific) restriction enzymes using InFusion HD cloning kit (Takara Bio). All plasmid constructs were verified by DNA sequencing.

Lentiviral CRISPR/Cas9 transfer vector was modified from lentiCRISPRv2 backbone plasmid (Zheng Lab, Addgene, 52961) (Sanjana et al., 2014). Using pCDH-hSYN-huLAMP1-mCherry as a template, a fragment containing P2A-mCherry-NLS was PCR amplified with Phusion High-Fidelity DNA polymerase (Thermo Scientific) and the following primers, P2A_Cher_for 5'TGACGATAAGGGATCCGGCGCAACAAACTTCTCTCTGCTGAAACAAGCCGGAGATGTCGAAGAGAATCCTGGACCGGTGAGCAAGGGCGAGGAG and NLS_Cher_rev 5'GTCTAGTTTTAACGCGTTTGGCAGCAGGCTTGTACAGCTCGTCCATGCC. From plentiCRISPRv2, WPRE-5'LTR was PCR amplified with Phusion High-Fidelity DNA polymerase (Thermo Scientific) and the following primers, NLS_WPRE_for 5'CGCGTTAAACTAGACTGAACGCGTTAAGTCGACAATCAAC and LTR_rev 5'CTGATCAGCGGGTTTAAACGGGCCCTGCTAGAGAT. Using InFusion HD cloning kit (Takara Bio), P2A-mCherry-NLS and WPRE-5'LTR fragments were cloned into plentiCRISPRv2 vector digested with FastDigest-*Bam*HI/-*Pme*I (Thermo Scientific). EF1a promoter driving the expression of *SpCas9*, was replaced with hSYN promoter by cloning it from pCDH-hSYN-T2A-EGFP to pLentiCRISPR-P2A-mCherry-NLS vector using FastDigest-*Nhe*I/*Xba*I restriction enzymes (Thermo Scientific) to obtain pLentiCRISPR-hSYN-Cas9-P2A-mCherryNLS vector. The sgRNA targeting *Ret,* 5'TCTATGGCGTCTACCGTACA, was designed with Optimized CRISPR design tool (http://crispr.mit.edu/) (Hsu et al., 2013) and scrambled (SCR) sgRNA sequence, 5'GCACTACCAGAGCTAACTCA, was from pCas-Scramble CRISPR Vector (OriGene, GE100003). As a pair of annealed oligos, each sgRNA sequence was cloned into pLentiCRISPR-hSYN-Cas9-P2A-mCherryNLS digested with FastDigest-*Bsm*BI, as described in (Sanjana et al., 2014). All plasmid constructs were verified by DNA sequencing.

**Production of lentivirus vectors**

Human HEK293T cells (ATCC, CRL-1573) were grown on 10-cm cell culture dishes in Dulbecco's Modified Eagle Medium (DMEM; Gibco, 12491-015; supplemented with 10% FBS (Gibco, 10500056) and 100 μg/mL normocin (Invivogen, ant-nr-1, ant-nr-2)). One day pre-transfection, cells were re-plated in cell culture medium containing 25 mM HEPES (N-2-hydroxyethylpiperazine-N-2-ethane sulfonic acid; Gibco, 15630080). Per vector, 8-12 10-cm dishes of cells were transfected by using linear polyethylenimine (PEI; Polysciences, 23966-2). Per dish, DNA mixture was prepared in 500 μL pre-warmed OptiMEM reduced serum medium (Gibco, 1985070); final concentration of 4 μg of transfer plasmid and 2 μg of each helper plasmids (pMDLg/pRRE, (Addgene, 12251), pRSV/REV (Addgene, 12253) and pMD2.G (Addgene, 12259)). PEI (1 μg/μL, in 1x PBS, pH 4.5) was added in 4:1 (PEI: DNA) v/w ratio, mixture was incubated at RT for 10 minutes, and then added to petri dishes, drop-wise. Dishes were incubated at 5% CO2, 37°C for 60-72 hours. Media was collected into Falcon tubes, centrifuged at 300x g for 5 minutes. After filtration with 0.45 μm filter, supernatant was transferred into Ultra-Clear centrifuge tubes (Beckman Couter, 344058) fitted into metal rotor tubes for AH-629 ultracentrifuge rotor (Beckman Coulter Inc, Brea, California, USA) and centrifuged at 120,000x g for 1.5 hours, at +4°C. Pellet was dried under the hood for 5-10 minutes, then re-suspended with Dulbecco's PBS and centrifuged for a minute at 17,000x g. Supernatant was collected, aliquoted, stored at -80°C.

**Supplementary Figures**

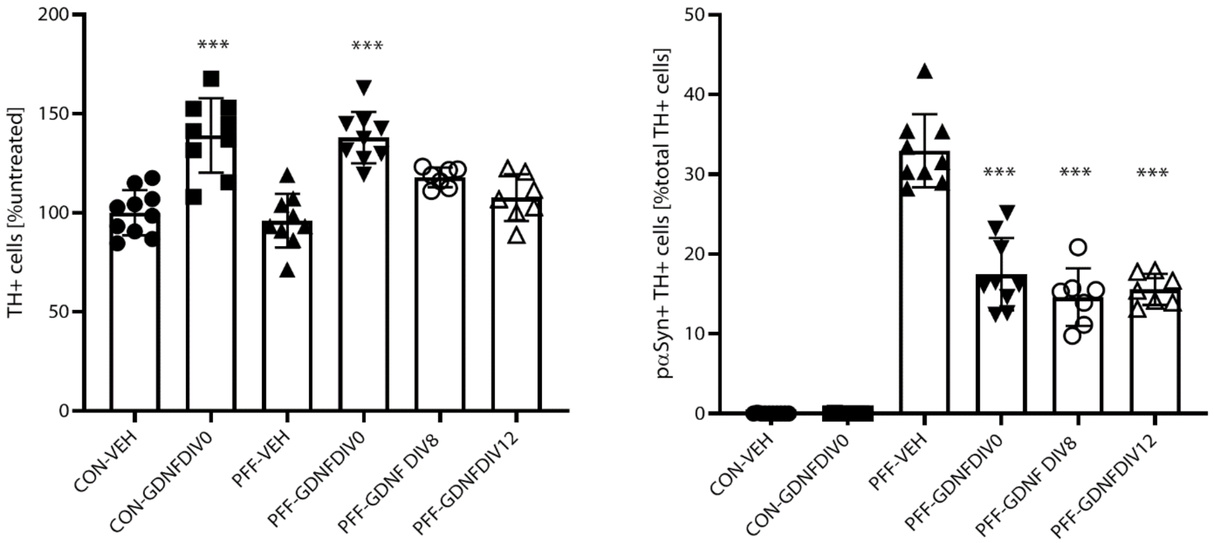


Fig S1. **GDNF treatment up to 4 days after PFF addition reduces the accumulation of α-synuclein aggregates in cultured dopamine neurons**. GDNF (50 ng/ml) was added to control (CON) or PFF treated wells at different time points (DIV0, DIV8, DIV12). In all cases, GDNF reduced amount of large, LB-like pSer129 α-synuclein aggregates in dopamine neurons (one way ANOVA and Holm-Sidak's test). As expected, when included from the beginning of the culture (DIV0), GDNF increased number of surviving TH-positive cells, but the effect was much less pronounced when the neurotrophic factor was added later, possibly, due to stabilization of dopamine cell number in the culture at earlier time points (Kruskal-Wallis and Dunn's tests). Data are mean ±SD, *p<0.05, ***p<0.001, vs CON-VEH, n=7-10 wells from two independent experiments.

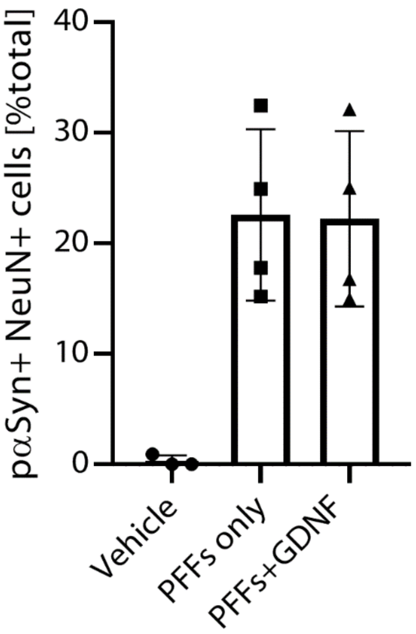


Fig S2. **GDNF does not prevent pαSyn accumulation in hippocampal neurons**. Cells were pretreated with GDNF (50 ng/ml) and subsequently with PFFs (PFFs+GDNF), just PFFs (PFFs only) or treated with PBS only (Vehicle). GDNF did not reduce the number of LB-like pSer129 α-synuclein aggregates in neurons identified by NeuN immunostaining when quantified 7 days after PFF treatment. Data are mean ±SD from 3-4 independent experiments.

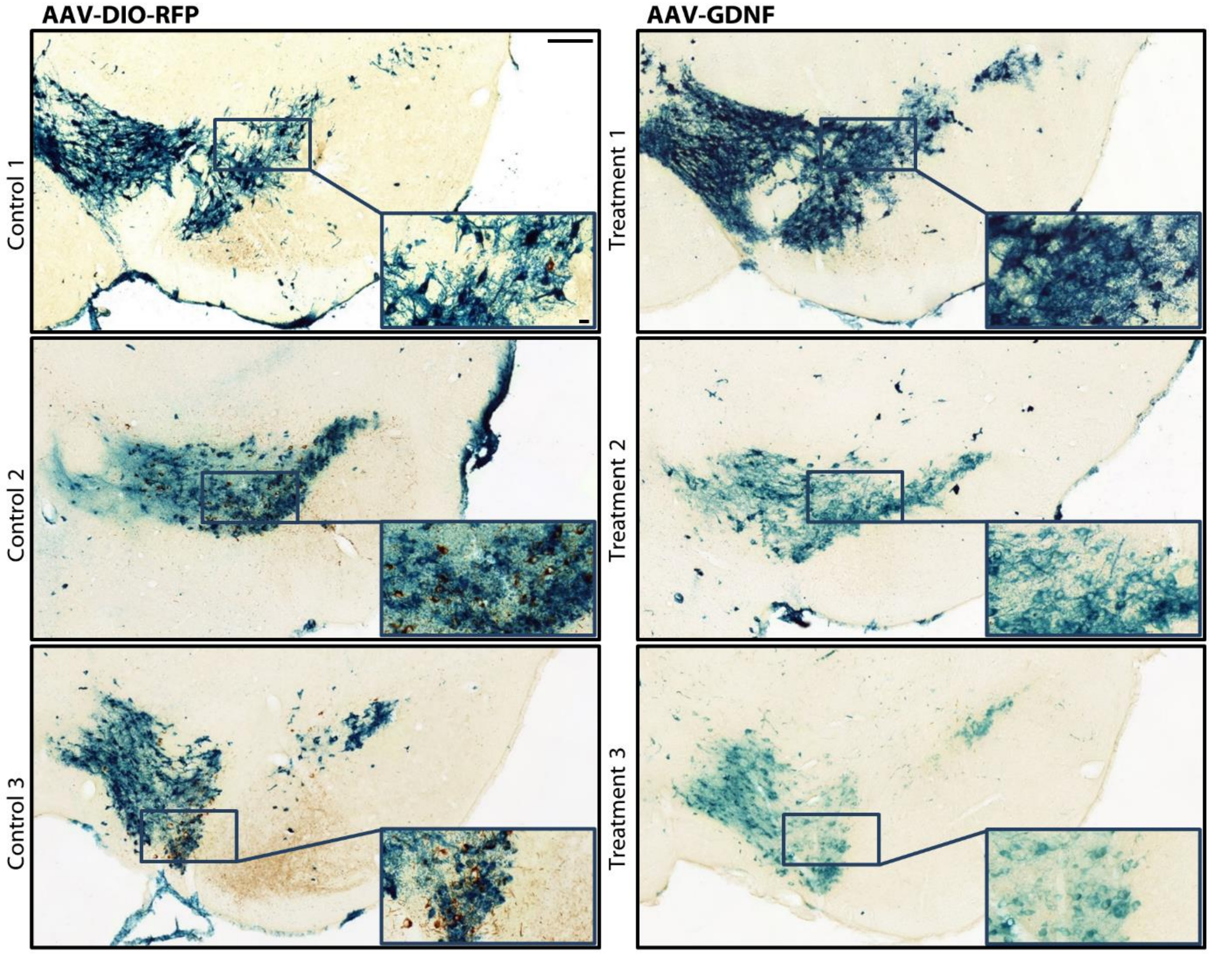


Fig S3. **Accumulation of LB-like pSer129 α-synuclein aggregates in the SN of PFF-injected mice with or without GDNF overexpression**. LB-like aggregates and LB-like neurites (both brown), were abundant in the SN of animals that received control vector (AAV-DIO-RFP, left). On the other hand, they were very scarce in the SN of animals that received the treatment vector (AAV-GDNF, right). The intensity of TH immunostaining (blue) varied in different animals of the treatment group (right). Scale bars, 200 µm and 20 µm, respectively. Each panel represents a different animal, n=6 (3 control + 3 treatment).
